# Adaptive laboratory evolution of *Escherichia coli* under acid stress

**DOI:** 10.1101/659706

**Authors:** Bin Du, Connor A. Olson, Anand V. Sastry, Xin Fang, Patrick V. Phaneuf, Ke Chen, Muyao Wu, Richard Szubin, Sibei Xu, Ye Gao, Ying Hefner, Adam M. Feist, Bernhard O. Palsson

## Abstract

The ability of *Escherichia coli* to tolerate acid stress is important for its survival and colonization in the human digestive tract. Here, we performed adaptive laboratory evolution of the laboratory strain *E. coli* K-12 MG1655 at pH 5.5 in glucose minimal medium. By 800 generations, six independent populations under evolution reached 18.0% higher growth rates than their starting strain at pH 5.5, while maintaining comparable growth rates to the starting strain at pH 7. We characterized the evolved strains to find that: (1) whole genome sequencing of isolated clones from each evolved population revealed mutations in *rpoC* appearing in 5 of 6 sequenced clones; (2) gene expression profiles revealed different strategies to mitigate acid stress, that are related to amino acid metabolism and energy production and conversion. Thus, a combination of adaptive laboratory evolution, genome resequencing, and expression profiling reveals, on a genome-scale, the strategies that *E. coli* deploys to mitigate acid stress.

## Introduction

As a commonly found enteric bacteria species in the human digestive tract, *Escherichia coli* is known to withstand various levels of acid stress [1–6]. For example, *E. coli* can survive several hours under pH 2 [1], which is within the range of the extremely acidic stomach (pH 1.5 to 3) that serves as a barrier for most bacteria [7]. Additionally, *E. coli* has been shown to grow under mild acid stress [4–6], which is typically found in the human intestinal tract [7, 8]. Such adaptability to low pH environments has raised wide interest in understanding the underlying mechanisms that protect *E. coli* from acid stress. Furthermore, studying the acid resistance mechanisms of *E. coli* has important implications in the food and health care industry. For example, treatment strategies can be developed to target specific acid resistance mechanisms in the case of a pathogenic *E. coli* infection.

The acid resistance mechanisms of *E. coli* have been studied extensively. To maintain intracellular pH homeostasis, *E. coli* has developed various strategies including cytoplasmic buffering [9], proton-consuming systems [10–13], adjustment of cellular metabolism [14, 15], and physiological responses [16–20]. The buffering capacity of the cytoplasm mainly comes from inorganic phosphates, amino acid side chains, polyphosphates, and polyamines [9]. The proton-consuming systems include four types of amino acid decarboxylase systems that function under different pH conditions and formate hydrogen lyase that is active under anaerobic conditions [21]. The metabolic responses under acid stress include the up-regulation of components in the electron transport chain and metabolism of sugar derivatives that have decreased acid production compared to glucose [14, 15]. The physiological responses include the activation of periplasmic chaperones HdeA and HdeB [16], adjustment of membrane lipid compositions [17, 18], and blockage of outer membrane porins [19, 20].

Adaptive laboratory evolution (ALE) is an important experimental approach for understanding the adaptive response of microorganisms to particular environments or after exposure to stresses [22]. During an ALE experiment, the microorganism is cultured under defined conditions for an extended period of time. ALE allows the selection of improved phenotypes, typically the growth rates, under certain growth environments. Furthermore, the advancement of next-generation sequencing technology makes it convenient to obtain the genotypes underlying the favorable traits over the course of evolution [23]. A previous study also investigated the adaptive evolution of *E. coli* under acid stress, where *E. coli* K-12 W3110 was evolved in a nutrient rich environment (LBK medium) buffered at pH range 4.6 - 4.8 for 2000 generations [24, 25]. Here, we are interested in the adaptive evolution of *E. coli* under acid stress in a nutrient limited environment, where glucose is the only carbon source.

In this study, we perform ALE on *E. coli* at pH 5.5 in glucose minimal medium. For the evolved strains, we use whole genome sequencing to identify genetic mutations that arise over the course of evolution. Additionally, to examine the change in gene activity after evolution, we perform RNA sequencing to characterize the gene expression profile of the evolved endpoints when growing under different pH conditions. We then identify the differentially expressed genes (DEGs) of the evolved endpoints at different pH conditions. We also uncover new cellular processes that emerge over the adaptive evolution under acid stress, using DEGs identified in the starting strain across pH as a reference.

## Methods

### Culture medium

The M9 glucose minimal medium was prepared by adding the following to Milli-Q water: 0.1 mM CaCl_2_, 0.2 mM MgSO_4_, 1× trace elements solution, 1× M9 salt solution, and 4 g/L D-glucose. Trace elements solution (4000×) was prepared in concentrated HCl with 27 g/L FeCl_3_ ***·*** 6H_2_O, 1.3 g/L ZnCl_2_, 2g/L CoCl_2_ ***·*** 6H_2_O, 2 g/L Na_2_MoO_4_ ***·*** 2H_2_O, 0.75 g/L CaCl_2_, 0.91 g/L CuCl_2_ ***·*** 2H_2_O, and 0.5 g/L H_3_BO_3_. M9 salt solution (10×) was prepared by dissolving 68 g/L Na_2_HPO_2_, 30 g/L KH_2_PO_4_, 5 g/L NaCl, and 10 g/L NH_4_Cl in Milli-Q water. It is worth mentioning that the concentration of MgSO_4_ is 10 times lower than used previously [26], as higher concentration of magnesium ion led to precipitation issues. To maintain the pH around 5.5 during cell culture, the culture medium was supplemented with 150 mM 2-(N-morpholino) ethanesulfonic acid (MES) buffer from a 500 mM stock prepared in Milli-Q water. After mixing all components of the medium, the pH was adjusted using 2 M H_2_SO_4_ and 4 M KOH. All stock solutions as well as the final medium were sterile filtered through a 0.22 µM PVDF membrane.

### Adaptive laboratory evolution process

Cultures were initiated from isolated colonies of an *Escherichia coli* K-12 MG1655 strain (ATCC 47076), which had previously been evolved for approximately 10^13^ cumulative cell divisions (CCDs) on M9 minimal medium supplemented with 4 g/L of glucose [26]. The cultures were first grown overnight and then placed in tubes on a platform that performed passage automatically. The working culture volume was 15 mL, and the culture temperature was maintained at 37 °C. The culture medium was magnetically stirred at 1100 rpm to ensure a well mixed and aerobic growth environment.

From the start of the culture to the next passage, on average 4 samples of 100 uL culture medium were taken and the optical density measurements at a wavelength of 600 nM (OD600) were performed in a spectrophotometer (Tecan Sunrise). To maintain the cells in exponential growth phase, the culture medium containing *E. coli* (100 uL) was passaged to a tube containing fresh medium when OD600 of the original medium approached 0.3. The growth rate was determined for each culture using a least-squares fit on ln(OD600) versus time. Growth trajectories were generated by fitting a monotonically increasing cubic-interpolating-spline to the calculated growth rate values versus CCDs, as described previously [26]. Glycerol stocks of the cultures were taken periodically by mixing 800 µL of sterile 50% glycerol with 800 µL of culture and storing at −80 °C.

Throughout the course of the evolution, the culture medium pH was constantly measured to ensure proper buffering (Table S1). Specifically, the pH values of the fresh medium and the culture medium before the next passage were measured. The culture medium was filtered through 0.22 µM membranes, and the pH was measured using a meter (Fisher Scientific Accumet AB15). Additionally, OD600 measurements of the culture medium were taken before the next passage to assess the possible effect of cell density on culture medium pH.

### Whole genome sequencing and analysis of genetic mutations

Genomic DNA was isolated using bead agitation as described previously in Marotz et al. [44]. Whole genome DNA sequencing libraries were generated using a Kapa HyperPlus library prep kit (Kapa Biosystems). The libraries were then run on an Illumina HiSeq 4000 platform with a HiSeq SBS kit and 150/150 paired-end reads. The raw DNA sequencing reads in fastq format were processed using the *breseq* computational pipeline v0.33 [28]. Specifically, the workflow includes quality control [45], alignment to the *E. coli* genome (NCBI accession NC_000913.3) to identify mutations and annotation of the mutations. It is worth mentioning that genomic DNA was extracted for individual clones taken at different CCDs of the evolution and at the end of the evolution. The mutations identified in the clones at the end of the evolution were reported and those found at earlier stages were used to track how different mutations emerge or disappear throughout the course of the evolution.

### RNA sequencing

RNA sequencing data were generated from cell cultures under exponential growth phase at pH 5.5 and pH 7. The culture conditions at pH 5.5 were the same as used in ALE experiments mentioned above. The culture conditions at pH 7 were the same as the regular M9 glucose minimal medium [26], with no additional modifications on the medium. Cells were stabilized with Qiagen RNA-protect Bacteria Reagent. Cell pellets were stored at −80 °C before RNA extraction. Then, frozen cell pellets were thawed and incubated with lysozyme, protease K, SuperaseIN, and 20% sodium dodecyl sulfate for 30 minutes at 4 °C. Total RNA was isolated and purified using Qiagen’s RNeasy Plus Mini Kit based on the manufacturer protocol. The total RNA quality was checked using the RNA 6000 Nano kit from Agilent Bioanalyzer. For gram-negative bacteria, ribosomal RNA was removed using Ribo-Zero rRNA removal kit from Epicentre. Single-end, strand-specific RNA-seq libraries were generated using KAPA RNA HyperPrep Kit from Kapa Biosystem. RNA-seq libraries were run on an illumina NextSeq platform using a 75 cycle mid-output kit.

### Analysis of differentially expressed genes (DEGs) on RNA sequencing data

Raw sequencing reads in fastq format were first mapped to the reference genome (NCBI accession NC_000913.3) using bowtie v1.2.2 [46]. The abundance of the transcript was obtained using the summarizeOverlaps function from the GenomicAlignments package in R [47]. From the transcript abundance, the DEGs between two conditions were identified through the DESeq2 package in Bioconductor [48]. The output for the DEGs include log_2_(fold change) and the corresponding p-values (FDR-adjusted). DEGs with log_2_(fold change) greater than 1 and p-value smaller than 0.01 were considered to be significantly changed between the two conditions compared.

### Enrichment analysis for cluster of orthologous group (COG) categories

The set of DEGs between two different conditions were annotated using COG categories. The hypergeometric test was then performed for the set of upregulated genes and downregulated genes, respectively. To calculate the enrichment of each COG category in the gene set, four values were obtained to perform the test: the total number of genes mapped in RNA-seq data, the number of genes in the current set, the number of genes with the current COG category out of all genes, the number of genes with the COG category out of the current gene set. The FDR correction was applied on the p-values of the COG categories in the gene set. COG category with corrected p-value smaller than 0.05 was considered enriched in the gene set.

### Data availability

The genomic sequence data have been deposited in the NCBI Sequence Read Archive (SRA) under BioProject ID PRJNA546056. The RNA sequencing data have been deposited in the NCBI SRA under BioProject ID PRJNA546062.

## Results

### Laboratory evolution and acid-adapted endpoint strains

We used wild-type *Escherichia coli* K-12 MG1655 that had been previously evolved on M9 glucose minimal medium as the starting strain for evolution and refer to it as GLU strain [26]. We used GLU as the starting strain to isolate changes due to adaptation to acid stress from those caused by adaptation to the culture medium. The genetic mutations of the GLU strain against *E. coli* K-12 MG1655 is also documented in ALEdb (aledb.org) under the experiment name GLU. Six independent cultures were established under pH 5.5 in glucose minimal medium, buffered with 150 mM MES (p*K*_a_ = 6.1) [27]. In addition, we lowered magnesium (Mg) concentration in the media to 0.2 mM to minimize precipitations. We refer to six acid-adapted strains as AA1 to AA6, respectively. To account for the possible effects due to changes in media composition, we also set up two independent cultures under pH 7 with 150 mM MES buffer (MES1, MES2) and lowered Mg concentration (LM1, LM2), respectively. All of the strains used in this study and their relationships can be found in Figure 1A.

**Figure 1.**
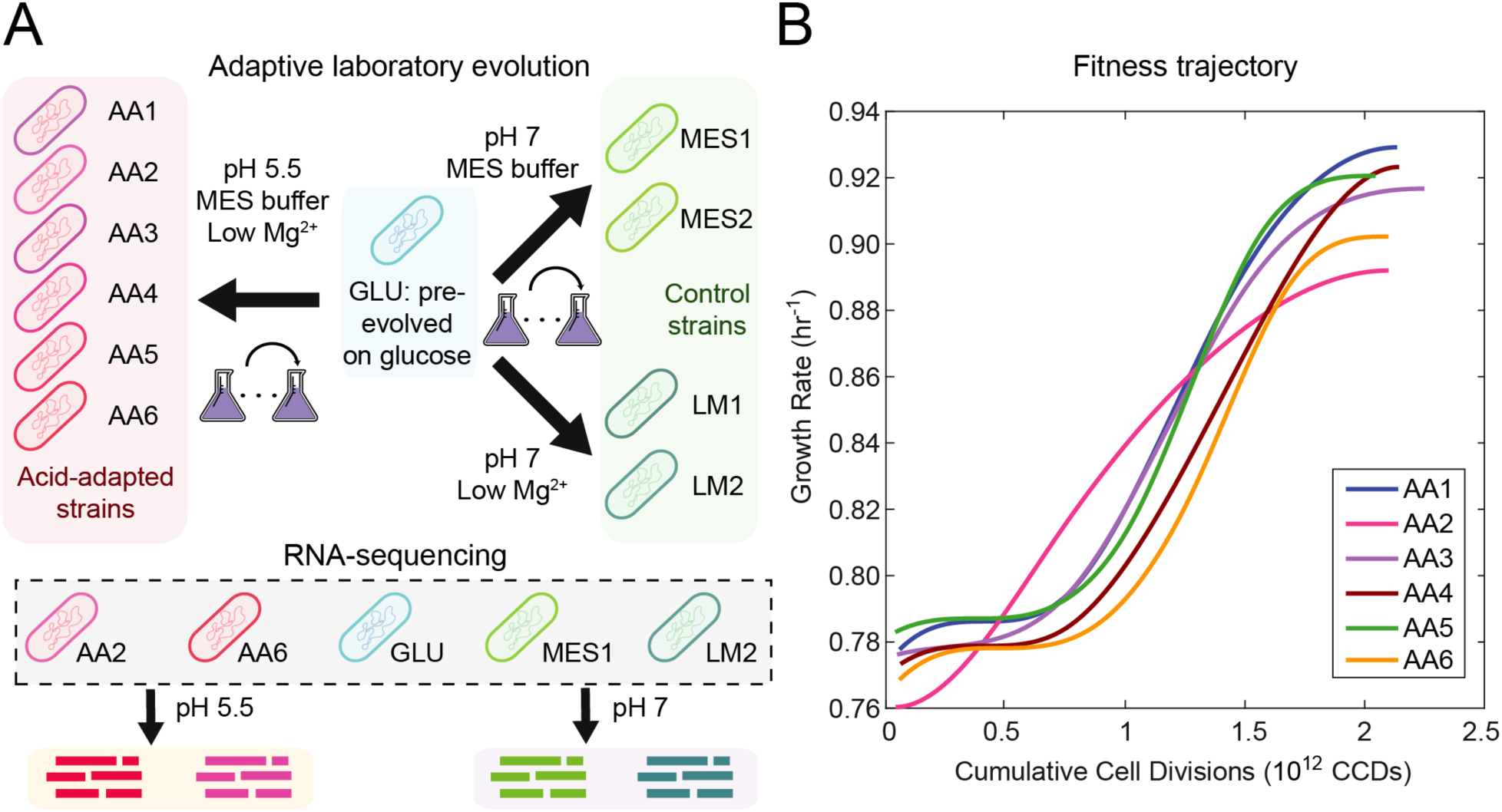
Adaptive laboratory evolution (ALE) of *E. coli* under acid stress. (A) Schematic for ALE process and strains used for RNA sequencing. Starting from the GLU strain, we performed ALE under pH 5.5 to obtain six acid-adapted (AA) strains and under pH 7 to obtain four control strains. Two control strains were adapted in glucose minimal medium with lowered magnesium concentration (LM) and two controls strains were adapted in glucose minmal medium with MES buffer (MES). Using RNA sequencing, we obtained the gene expression profiles of five selected strains under pH 5.5 and 7. (B) Smoothed fitness trajectories of six acid-adapted strains (raw data in Supplementary Figure S1). We show here the change of growth rate over cumulative cell divisions through the evolution process. The average growth rate improvement is 18%.

We performed ALE using an automated system, which tracked culture growth rates and passed the cells to fresh media when OD600 measurements reached 0.3 to ensure selection at exponential-phase growth. Additionally, we periodically measured the pH of the clean media and recently passaged cultures to ensure proper buffering (Table S1). The culture pH remained relatively stable for strains evolved in MES buffer under pH 5.5 and 7. For strains evolved under lowered Mg concentration and with only phosphate buffers (p*K*_a_ = 7.2) in glucose minimal medium, the culture pH dropped significantly at the end of the culture, likely due to the secretion of organic acids during growth. The laboratory evolution process lasted 35 days for strains AA1 to AA6 under pH 5.5, corresponding to 800 generations and 2.1 × 10^12^ CCDs. The fitness trajectories of the evolved strains are shown in Figure 1B. We observed the growth rates to continuously improve over CCDs and approach stable values at the end of the evolution. Overall, we found the evolved endpoints to have an average of 18.0% improvement in growth rate (from 0.77 ± 0.01 hour^−1^ to 0.91 ± 0.01 hour^−1^) over their starting strain.

To evaluate the fitness of acid-adapted strains against the starting GLU strain, we obtained the growth rates of the strains under pH 5.5 and 7 in a separate experiment. We found the acid-adapted strains to have increased fitness under pH 5.5, with growth rate at 0.83 ± 0.01 hour^−1^ compared to the growth rate at 0.67 ± 0.02 hour^−1^ of the GLU strain. We also found the growth rates under pH 7 for acid-adapted strains to be 1.00 ± 0.01 hour^−1^, slightly higher than that of the GLU strain at 0.94 ± 0.03 hour^−1^.

### Genetic mutations of the evolved strains

To understand the genetic basis of the observed phenotypic change, we performed whole genome sequencing on individual clones picked from acid-adapted strains AA1 to AA6, as well as the control strains MES1, MES2, LM1, and LM2. We identified the genetic mutations of the evolved strains by comparing them to the reference genome using *breseq* computational pipeline v0.33 (Methods) [28]. We reported the converged mutations of acid-adapted strains in Table 1. Converged mutations are mutations on the same gene identified across multiple strains from independent cultures. All mutations identified in acid-adapted strains and those of control strains can be found in Table S2 and S3.

**Table 1.** Converged mutations identified in the clones of acid-adapted strains under pH 5.5.

| Gene | Mutation | Protein change | Flask number | AA1 | AA2 | AA3 | AA4 | AA5 | AA6 |
| --- | --- | --- | --- | --- | --- | --- | --- | --- | --- |
| *rho* | C→A | R102S (CGC→AGC) | 87 | X |  |  |  |  |  |
| *rho* | C→T | R102C (CGC→TGC) | **111** |  |  |  |  |  | X |
| *rpoC* | C→A | A397E (GCG→GAG) | **111** |  | X |  |  |  |  |
| *rpoC* | G→C | G444A (GGT→GCT) | 88, **118** |  |  | X |  |  |  |
| *rpoC* | Δ7 bp | coding (4106 - 4112/4224 nt) | **113** |  |  |  | X |  |  |
| *rpoC* | Δ1 bp | coding (4111/4224 nt) | **111** |  |  |  |  | X |  |
| *rpoC* | C→T | S539F (TCT→TTT) | **111** |  |  |  |  |  | X |
| *nagA* | G→C | S90* (TCA→TGA) | 84 |  |  |  | X |  |  |
| *nagA* | C→T | R149H (CGT→CAT) | 83 |  |  |  |  |  | X |

Flask number stands for the number of cell passages during the evolution. The numbers in bold represent the final flask number (thus the endpoint) in the evolution process of the specific acid-adapted strain. The non-bold numbers are intermediate flasks during the evolution where samples were selected for whole genome sequencing.

Overall, we found a total of 22 mutations in all acid-adapted strains, including those of clones picked from the endpoints (Table 1 and S2 bold flask number) and midpoints of the evolution (Table 1 and S2 non-bold flask number). Notably, we observed mutations in *rpoC* to appear in 5 of the 6 endpoint clones. The *rpoC* gene encodes a subunit of RNA polymerase, which is known to act as a global regulator for gene expressions [29, 30]. The mutations on *rpoC* include both single nucleotide polymorphisms (SNPs) and deletions. We found these mutations to be located on the interaction interfaces of the protein product. Specifically, mutation with protein change A397E (Table 1) is located at the exit gate of the newly synthesized RNA strand. The region with mutation G444A interacts with rpoB subunit and the region with mutation S539F interacts with rpoA subunit (Table 1). The two deletion mutations are located at the interaction interface with rpoZ subunit (Table 1). Mutations in *rpoC* in *E. coli* have been found in several previous ALE experiments, covering a variety of experimental conditions or perturbations, e.g. high temperature, alternating substrate, gene knockouts [31–33]. Several studies have suggested mutations in *rpoC* to be mainly associated with improvement in metabolic efficiency and growth rate [34–36].

The other converged mutations are found in *rho* (transcription regulation) and nagA (metabolism of N-acetyl-D-glucosamine) (Table 1). The mutations in these two genes are all SNPs. Unlike *rpoC* where mutations are found in endpoint clones, mutations in *rho* appear in the midpoint clone of strain AA1 and endpoint clone of strain AA6. Mutations in *nagA* appear in midpoint clones of strain AA4 and AA6 and are not found in the endpoint clones of these two strains. For the rest of the mutations observed in acid-adapted strains, each of them appear only in a single strain (Table S2). These mutations appeared in both the coding regions and intergenic regions. The types of mutations include SNPs, deletions, and insertion elements.

We found distinct patterns when examining mutations in control strains evolved under different conditions (Table S3). We discovered SNPs on *oxyR* gene to be the converged mutations in strains LM1 and LM2. On the other hand, the converged mutations for strains MES1 and MES2 are found in the intergenic region of *ilvL* and *ilvX*. Notably, the exact same mutation between *ilvL* and *ilvX* is also found in the endpoint clone of acid-adapted strain AA5 (Table S2), confirming the possible effect of the MES buffer during the evolution process.

### Differential gene expression of the evolved endpoints at different pH conditions

To understand how the mutations can affect gene products in evolved strains, we used RNA sequencing to examine the gene expressions of ALE endpoints. We selected two acid-adapted strains for RNA-seq, strain AA2 which has a single mutation in *rpoC* and strain AA6 which has the most number of mutations among all AA strains (Table S2). We selected strains MES1 and LM2 for different control conditions. We performed RNA sequencing and obtained gene expression profiles of the selected strains as well as the starting GLU strain grown under pH 7 and pH 5.5 (Figure 1A; Methods). We then analyzed the expression profiles on the level of individual genes and their related cellular processes. Specifically, we performed statistical tests to identify the DEGs of the same strain growing under pH 5.5 compared to pH 7 (Methods).

To ensure the DEGs identified for acid-adapted strains across pH are only due to the effect of adaptive evolution under acid stress, we first need to understand the response to acid stress of the starting strain and also control for possible variations in culture medium during the evolution process. Therefore, we examined the DEGs across pH for the GLU strain, as well as the MES1 strain and LM2 strain to account for the possible effects due to MES buffer and lowered magnesium concentration. We found significant overlap of upregulated genes involved in cell wall/membrane biogenesis and translation processes among the acid-adapted strains and the control strains, implicating these two cellular processes as the common acid resistance mechanisms in *E. coli*.

We then examined DEGs in the acid-adapted strains under pH 5.5 compared to pH 7, after removing DEGs also found in GLU and the control strains. We found 183 genes to be differentially expressed for strain AA2 (111 upregulated and 72 downregulated) and 40 genes for strain AA6 (38 upregulated and 2 downregulated) (Table S4 and S5). Of those genes, we found 14 upregulated genes that appeared in both acid-adapted strains (Figure 2A). Based on COG annotation [37], the 14 genes were found to be mainly involved in energy production and conversion (e.g., TCA cycle, respiratory chain, ATP synthase) and amino acid transport and metabolism (e.g., biosynthesis of glutamate).

**Figure 2.**
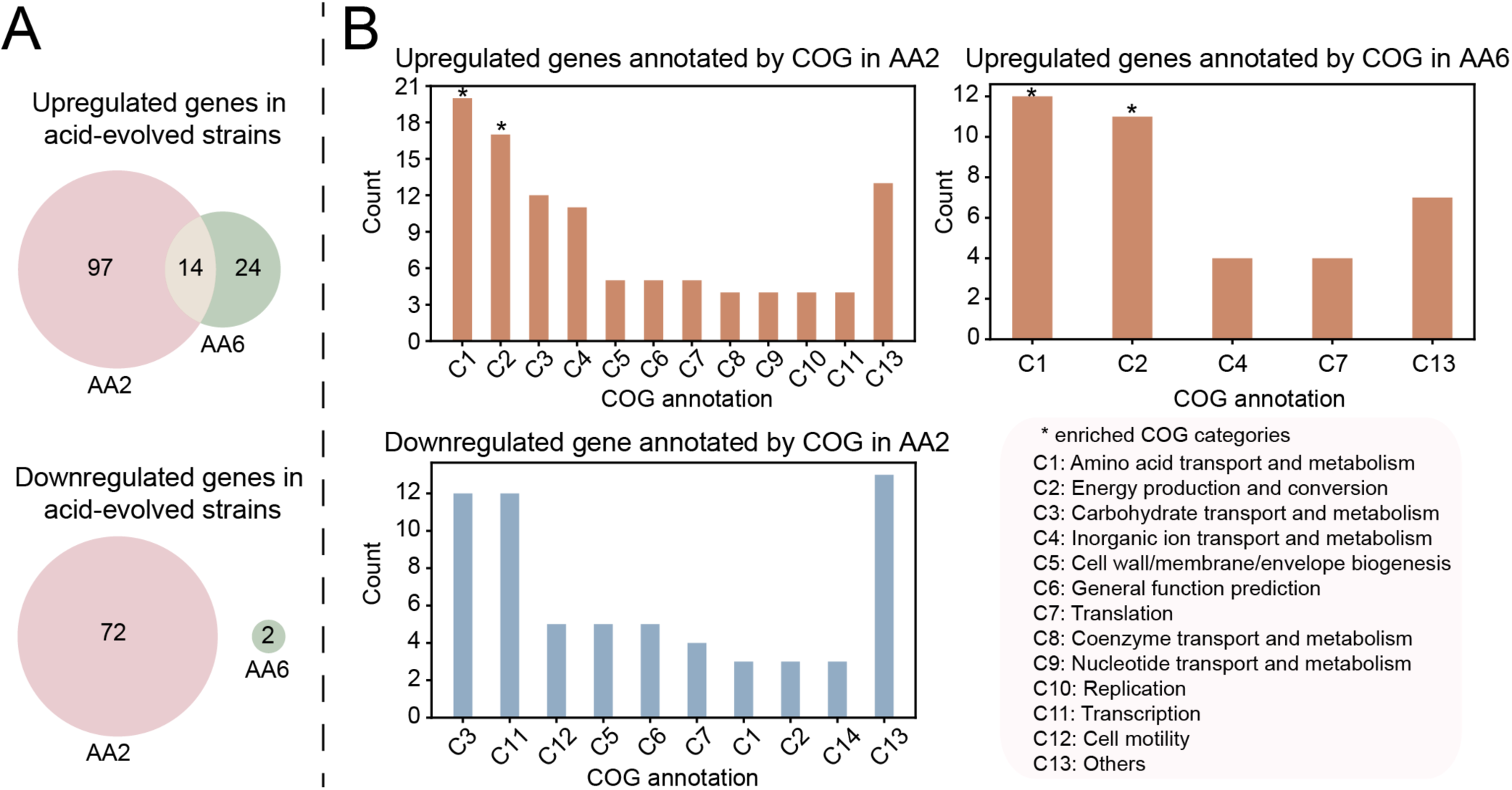
Differentially expressed genes (DEGs) of acid-adapted (AA) strains at different pH conditions. The DEGs are calculated for the same strain by comparing its gene expression profiles when growing under pH 5.5 and pH 7. (A) Number of upregulated (top panel) and downregulated genes (bottom panel) in acid-adapted strains AA2 and AA6. We found AA2 and AA6 to share 14 upregulated genes. (B) Cluster of orthologous group (COG) categories in the upregulated and downregulated genes in acid-adapted strains. The asterisk sign on top of the bar means that the COG category is enriched, as calculated using a hypergeometric test (Methods). We did not show the COG categories for downregulated genes in strain AA6 since there are only two of them. The upregulated and downregulated genes for strains AA2 and AA6 are listed in Table S4 and S5.

We next examined the specific COG categories of the DEGs across pH in each acid-adapted strain. For strain AA2, the upregulated genes are associated with more than ten COG categories (Figure 2B), with the largest number of genes on amino acid transport and metabolism (e.g., biosynthesis of histidine, threonine), energy production and conversion (e.g., nitrite/nitrate reductase, succinate dehydrogenase, TCA cycle, etc.), carbohydrate transport and metabolism (e.g., glycolysis), and inorganic ion transport and metabolism (e.g., transport of iron, zinc, nitrite, nitrate). Among the processes identified, the upregulated genes are enriched in amino acid transport and metabolism and energy production and conversion based on a hypergeometric test (Figure 2B stars) (Methods). The downregulated genes of strain AA2 are found mostly in carbohydrate transport and metabolism (e.g., secondary carbon sources such as xylose, arabinose) and transcription. No COG categories are found to be enriched in those downregulated genes. For the other acid-adapted strain, AA6, the upregulated genes are mainly enriched in amino acid transport and metabolism (e.g., biosynthesis of glutamate, arginine) and energy production and conversion (e.g., TCA cycle, respiratory chain, ATP synthase) based on a hypergeometric test (Figure 2B stars). Again, no COG categories are enriched in the downregulated genes in strain AA6.

Overall, we observed the DEGs to be enriched in similar general COG categories for two acid-adapted strains, AA2 and AA6. However, the specific underlying cellular processes still differ, indicating different strategies developed by *E. coli* over the course of evolution under acid stress. Such differences cover different processes that are upregulated in amino acid biosynthesis (e.g., histidine for AA2 and glutamate for AA6) and energy production and conversion (e.g., anaerobic respiration found in AA2 but not in AA6) between two acid-adapted strains. Additionally, we found the downregulated genes to be involved in different COG processes between these two strains.

## Discussion

In this work, we used ALE to study the adaptation of *E. coli* K-12 MG1655 under acid stress in glucose minimal medium. Using whole genome sequencing, we identified mutations on *rpoC, rho*, and *nagA* to be the converged mutations in acid-adapted strains. We then used RNA sequencing to examine the gene expression profiles of acid-adapted strains across pH and compared them to those of the starting GLU strain and the control strains. Through analysis of DEGs, we identified cellular processes acquired by *E. coli* through the adaptive evolution under acid stress.

We found five of the six acid-adapted strains to have mutations in *rpoC*, which functions as a subunit of RNA polymerase. The specific mutations include substitutions that change the encoded amino acids (AA2, AA3, and AA6) and deletions in the coding region that lead to shifts in reading frame (AA4 and AA5). Based on the Pfam database, the substitutions occurred in the protein domains that contain the active site and the pore region that allows the entrance of nucleotides to the active site [38–40]. It is worth noting that the deletions occurred at the end of *rpoC* gene (base pair 4106 and 4111 out of 4224 base pairs) and likely did not result in significant disruption of the gene function. A previous study on the evolution of *E. coli* under acid stress by Harden et al. also observed missense mutations in subunits of RNA polymerase (*rpoBCD*) for all of the acid-adapted strains [24]. The authors in that study proposed several mechanisms to explain how mutations on the RNA polymerase complex might enhance fitness under acid stress. Here, however, we consider the mutations on *rpoC* to be associated with inducing faster growth rather than acid resistance. A comprehensive analysis on the 278 gene expression datasets of *E. coli* across diverse conditions has revealed that mutations on genes related to RNA polymerase typically lead to improved growth rate and reduced stress-related gene expression [41].

Other mutations found cover a range of cellular processes. However, none of the processes are directly related to the commonly known acid resistance mechanisms. A previous study by Harden et al. identified mutations in genes related to the amino acid decarboxylase systems, and these mutations result in loss or downregulation of amino-acid decarboxylase activities [24, 25]. The different mutations observed from the two studies are likely due to the different culture media in which the evolution took place. The culture medium used by Harden et al. is LBK medium, which is rich in amino acids. The activation of the amino acid decarboxylase systems requires the presence of amino acids in the medium [10–13]. According to Harden et al., amino acid decarboxylase systems protect *E. coli* from acid stress upon early exposure to the acidic environment, but incur fitness costs over the long term, where *E. coli* has developed other strategies to maintain the non-stress physiology. In our study, the culture medium is glucose minimal medium, where the sole carbon source is glucose. Therefore, the amino acid decarboxylase systems are never activated under this condition. Rather, we see mutations in genes that might be related to general cellular responses under stress conditions, e.g., transcription regulation (*rho*) and cellular physiology (*csgD/csgB, yiaA*).

For the two acid-adapted strains with gene expression profiles available, they share several common general COG categories for upregulated genes under acid stress. However, besides sharing 14 DEGs in processes such as TCA cycle, respiratory chain, ATP synthase, and glutamate biosynthesis, the two strains have a number of DEGs with different cellular functions (Figure 2). Both strains have SNPs in the *rpoC* gene, but in different protein domains according to the Pfam database mentioned earlier. Additionally, mutations on other genes in strain AA6 can possibly contribute to the different strategies used by *E. coli* under acid stress to adjust the level of gene transcripts. Similarly, the follow-up study on the work by Harden et al. also observed different patterns of gene expression across four acid-adapted strains [25]. These two studies together demonstrate that regardless of the level of acid stress (pH 4.6 - 4.8 by Harden et al. and pH 5.5 in this study) and nutrient availability (LBK medium and glucose minimal medium), the evolutionary pressure can drive *E. coli* to develop different strategies against acid stress.

Overall, we studied the adaptive evolution of *E. coli* under acid stress, linking the improved phenotype to the underlying genotypes and levels of gene expression. The study here provides a novel perspective on acid resistance mechanisms, as the commonly known acid resistance systems depend on rich medium or specific amino acids [42, 43]. In addition to the analysis of genetic mutations and DEGs, further analysis can be performed to understand the change in regulatory actions using a recently developed approach [41]. Such analysis can be helpful in understanding the response to acid stress at the level of transcriptional regulation and revealing potential drivers behind the global adjustment of cellular response against acid stress.

## Author statements

### Authors and contributors

BOP acquired the funding for this study. BOP, AMF and BD conceptualize the research goals and aims. BOP, AMF, BD and CAO designed the adaptive laboratory experiments. CAO, RS, SX, YG and YH performed the experiments, including adaptive laboratory evolution, whole genome sequencing and RNA sequencing. BD, AVS, XF, PVP, KC, MW performed the data analysis. BD, CAO, RS and BOP wrote the manuscript. All authors read and approved the final manuscript.

## Acknowledgements

We would like to thank Laurence Yang for valuable discussions.

