## Supplemental tables and figures for "Adaptive laboratory evolution of *Escherichia coli* under acid stress"

Table S1. Periodic pH measurements of culture media over the course of evolution.

| Day  Condition | 3 | 12 | 25 | 27 | 35 |
| --- | --- | --- | --- | --- | --- |
| pH 5.5 MES start | 5.55 | 5.45 |  | 5.51 | 5.52 |
| pH 5.5 MES end | 5.38 | 4.86 (0.26), 4.79 (0.39) |  | 5.14 (0.28) | 5.16 (0.37) |
| pH 7 MES start | 7.00 | 7.04 |  |  |  |
| pH 7 MES end | 6.84 | 6.54 (0.39) | 6.55 (0.26) |  |  |
| pH 7 low Mg start | 7.07 | 7.10 |  | 6.98 | 7.07 |
| pH 7 low Mg end | 6.02 | 5.76 (0.35) | 4.98 (0.28) | 5.86 (0.37) | 5.40 (0.49) |

Aliquots were taken from both clean media (start) and those already used for cell culture (end). Days for sampling were chosen at random, and the used media were obtained from tubes containing cells ready for next passage. The OD600 measurements of the culture media are shown in parenthesis.

Table S2. All mutations identified in the clones of acid-evolved strains under pH 5.5.

| Gene | Mutation | Protein change | Flask number | AA1 | AA2 | AA3 | AA4 | AA5 | AA6 |
| --- | --- | --- | --- | --- | --- | --- | --- | --- | --- |
| *yejH* | T→G | D105E (GAT→GAG) | 56 | X |  |  |  |  |  |
| *rho* | C→A | R102S (CGC→AGC) | 87 | X |  |  |  |  |  |
| *tam* | IS4 | coding (5-16/759 nt) | **114** | X |  |  |  |  |  |
| *ilvL* | Δ2 bp | coding (33-34/99 nt) | **114** | X |  |  |  |  |  |
| *rpoC* | C→A | A397E (GCG→GAG) | **111** |  | X |  |  |  |  |
| *rpoC* | G→C | G444A (GGT→GCT) | 88, **118** |  |  | X |  |  |  |
| *yraJ* | IS2 | coding (1893-1896/2517 nt) | 88 |  |  | X |  |  |  |
| *yiaA* | IS2 | coding (16-20/438 nt) | **118** |  |  | X |  |  |  |
| *nagA* | G→C | S90* (TCA→TGA) | 84 |  |  |  | X |  |  |
| *ycjX* | G→T | G207G (GGG→GGT) | 84 |  |  |  | X |  |  |
| *rpoC* | Δ7 bp | coding (4106-4112/4224 nt) | **113** |  |  |  | X |  |  |
| *ilvL/ilvX* | T→G | intergenic (+49/-38) | 69 |  |  |  |  | X |  |
| *yjcB* | C→T | R12H (CGC→CAC) | 69 |  |  |  |  | X |  |
| *rpoC* | Δ1 bp | coding (4111/4224 nt) | **111** |  |  |  |  | X |  |
| *metF/katG* | C→T | intergenic (+275/-54) | **111** |  |  |  |  | X |  |
| *csgD/csgB* | T→C | intergenic (-194/‑561) | **111** |  |  |  |  | X |  |
| *ydhY/ydhZ* | IS5 | intergenic (-215/+237) | 83, **111** |  |  |  |  |  | X |
| *nagA* | C→T | R149H (CGT→CAT) | 83 |  |  |  |  |  | X |
| *phoQ* | G→C | P208R (CCG→CGG) | 83 |  |  |  |  |  | X |
| *rho* | C→T | R102C (CGC→TGC) | **111** |  |  |  |  |  | X |
| *rpoC* | C→T | S539F (TCT→TTT) | **111** |  |  |  |  |  | X |
| *ycgJ* | IS2 | coding (1-5/369 nt) | **111** |  |  |  |  |  | X |

Flask number stands for the number of cell passages during the evolution. The numbers in bold represent the final flask number (thus the endpoint) in the evolution process of the specific acid-evolved strain. The non-bold numbers are intermediate flasks during the evolution where samples were selected for whole genome sequencing.

Table S3. Mutations identified in the clones of control strains evolved under lowered magnesium concentration or in MES buffer

| Gene | Mutation | Protein change | Flask number | LM1 | LM2 | MES1 | MES2 |
| --- | --- | --- | --- | --- | --- | --- | --- |
| *oxyR* | T→G | C199G (TGT→GGT) | **122** | X |  |  |  |
| *dsbG/ahpC* | IS5 | intergenic (-311/‑58) | **122** | X |  |  |  |
| *oxyR* | C→A | A204E (GCA→GAA) | **109** |  | X |  |  |
| *alaC/ypdA* | C→A | intergenic (-281/‑95) | **109** |  | X |  |  |
| *ilvL/ilvX* | T→G | intergenic (+49/-38) | **53**, **47** |  |  | X | X |
| *cbl* | G→T | Q225K (CAG→AAG) | **47** |  |  |  | X |
| *potG* | C→T | T107T (ACC→ACT) | **47** |  |  |  | X |
| *casA* | C→T | P60P (CCG→CCA) | **47** |  |  |  | X |

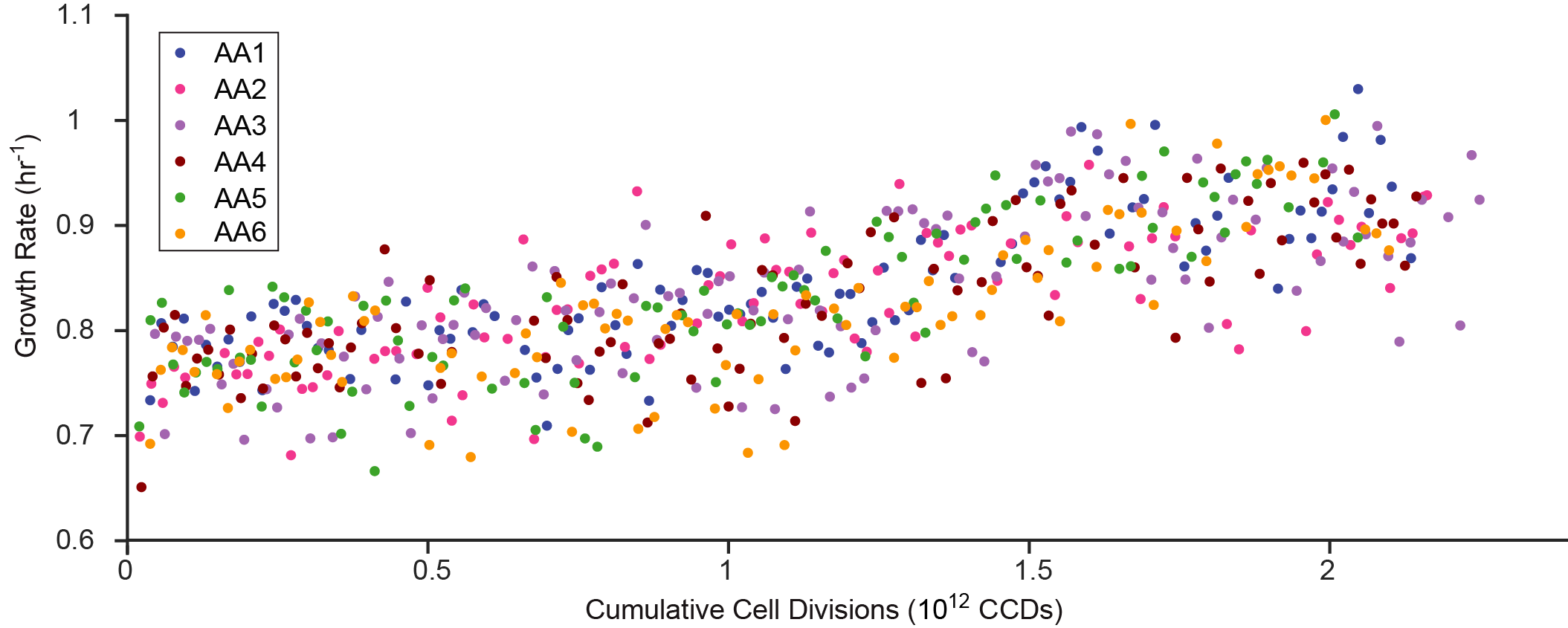

Figure S1. Raw data points for the evolution process of six acid-adapted (AA) strains.

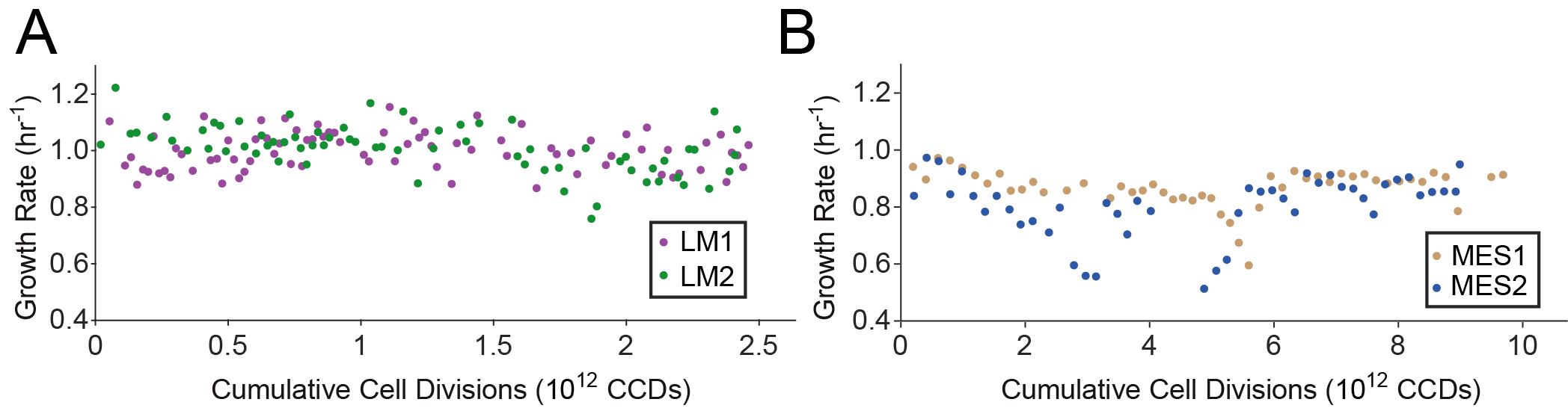

Figure S2. Growth rate versus cumulative cell divisions through the evolution process for control strains. (A) Control strains evolved in lowered magnesium concentration. (B) Control strains evolved in MES buffer.
